# Clinical and MRI changes of *puborectalis* and *iliococcygeus* after a short period of intensive pelvic floor muscles training with or without instrumentation

**DOI:** 10.1101/248823

**Authors:** Frédéric Dierick, Ekaterina Galtsova, Clara Lauer, Fabien Buisseret, Anne-France Bouché, Laurent Martin

## Abstract

**Purpose:** This study evaluates the impact of a 3-week period of intensive pelvic floor muscles training (PFMT), with or without instrumentation, on clinical and static magnetic resonance imaging (MRI) changes of *puborectalis* (PR) and *iliococcygeus* (IL) muscles.

**Methods:** 24 healthy young 5 women were enrolled in the study and 17 achieved the 9 sessions of 30 minutes training exercises and conducted all assessments. Participants were randomly assigned in two training groups: voluntary contractions combined with hypopressive exercises (HYPO) or biofeedback exercises combined with transvaginal electrical stimulations (ELEC). Clinical and T2-weighted MRI assessments io were realized before and after training.

**Results:** Modified Oxford Grading System (MOGS) scores for left PR and perineal body significantly increased in the two groups (*p*=0.039, *p*=0.008), but MOGS score for right PR significantly increased only in HYPO (*p*=0.020). Muscle volumes of right and left IL significantly decreased (*p*=0.040, *p*=0.045) after training as well as signal i5 intensities of right and left PR (*p*=0.040, *p*=0.021) and thickness of right and left IL at mid-vagina location (*p*=0.012, *p*=0.011).

**Conclusions:** A short period of intensive PFMT induces clinical and morphological changes in PFMs at rest suggesting a decrease in IL volume and adipose content of PR. Given the difference in cost and accessibility between HYPO and ELEC approaches, PFMT should be based primarily on non-instrumented exercises.

## 1 Introduction

Pelvic floor dysfunction (PFD), in particular stress urinary incontinence (UI) caused by structural defects in connective tissue and muscles that support pelvic organs (Petros, 2010; Memon and Handa, 2013), is a common and distressing condition affecting young and middle-aged women (Kim et al, 2003; Lose, 2005; Robinson and Cardozowan, 2014; Schreiber Pedersen et al, 2017). Several randomized controlled trials have demonstrated that pelvic floor muscles training (PFMT) is effective to reduce stress and any type of UI in women, therefore it is recommended as a first-line conservative therapy (Dumoulin et al, 2014, 2015). Nowadays, PFMT can be achieved with instrumentation, using vaginal/ anal probe electrode delivering biofeedback exercises or electrical stimulations, and weighted vaginal cones or other resistance devices. It can be also achieved without any instrumentation, using direct voluntary contractions, also known as Kegel’s exercises (Harvey, 2003), or indirect contractions with *transversus abdominis* muscle activation through hypopressive exercises (Caufriez, 1997; Resende et al, 2012). Previous studies on women with PFD show that PFMT without the use of electrical stimulation (Bø et al, 1999) or biofeedback (Fitz et al, 2012) is more efficient that with it, but this is controversial (Castro et al, 2008; Ibrahim et al, 2015). Uncertainty about which of these strategies are most effective in training women is one of the key clinical questions which needs to be prioritized because training using instrumentation is more costly than without. Therefore, further studies are needed to assess PFMT strategies efficacy.

The most common type of involuntary leakage of urine is stress urinary incontinence (SUI), a worldwide problem that affects the quality of life of millions of women since is typically experienced when coughing, sneezing, laughing, or exercising. In SUI, urine leakage is explained by a lack of closure pressure in the urethra during exertion, that could be attributed to anatomic changes in the bladder and urethra and/or PFM, endopelvic fascia and connective tissue supports (Petros and Ulmsten, 1990; Tunn et al, 2006; Hay-Smith et al, 2011). From a physiological point of view, we believe that it is relevant to study the impact of PFMT in young asymptomatic women with “non-pathological” *levator ani* (LA) muscle signals that are preserved from the consequences of parity or loco-regional surgery. However, to the best of our knowledge, no study has investigated clinical and morphological changes in LA muscle subdivisions before and after PFMT in young asymptomatic nulliparous women, even though a recent study reported a high prevalence of 20% for involuntary loss of urine in a group of 159 young presumably healthy women aged 18–30 years (van Breda et al, 2015).

The purpose of this randomized study was to evaluate the impact of a 3-week period of intensive PFMT, with or without instrumentation based on biofeedback and electrical stimulation, on clinical and static MRI changes of *puborectalis* (PR) and *iliococcygeus* (IL) muscles. The protocol without biofeedback and electrical stimulation was entirely realized without any instrumentation and based on voluntary PFM contractions combined with hypopressive exercises.

## 2 Methods

### 2.1 Study participants and PFM exercises

A convenience sample of 24 healthy young nulliparous women (age: 22.9±1.6 yrs, weight: 61.5±6.9 *kg*, height: 167.7±7.9 *cm*, body mass index (BMI): 21.8±1.5 *kg m*^‒2^), was recruited through advertisement and snowball sampling at our physiotherapy department. Before clinical and MRI assessments, all participants were questioned about PFD including symptoms of urinary and fecal continence, using a validated french version (de Tayrac et al, 2007) of Pelvic Floor Distress Inventory (PFDI-20) and Pelvic Floor Impact Questionnaire (PFIQ-7) (Barber et al, 2005). Additionally, we asked them if they were aware of their PFM function and if they known or already practiced PFM exercises. To be included in the study, participant had to have no MRI incompatibility (intrauterine device, pregnancy) or have practiced PFM exercises within the last year. Exclusion criteria were: pelvic floor troubles, diarrhea or chronic constipation, intensive sports practices, claustrophobia, neuroleptic or antidepressant medication, abortion. Inclusion criteria were: BMI<30 *kg m*^‒2^, intact perineum, age<30 *yrs*, nulliparous European women.

Participants were randomly assigned to two different PFM training groups. A first group realized voluntary PFM contractions combined with hypopressive exercises (HYPO) and the second followed biofeedback exercises combined with PFM electrical stimulations (ELEC). Participants were matched according to age (±1 *yr*) and BMI (±0.5 *kg m*^‒2^). The enrollment period was approximately 10–11 months. Participants of the two groups followed a total of 270 minutes (9 sessions of 30 minutes, 3 sessions/week during 3 weeks) PFMT, which are usually prescribed and reimbursed by Belgian health-insurance mutuals for uro-gynecologycal physiotherapy rehabilitation for the treatment of female UI.

In HYPO group, hypopressive abdominal exercises were instructed to the participants by 2 physiotherapists (E.G. and C.L.) and performed individually under their supervision, respecting the basic sequence proposed by Caufriez (1997): (1) slow and deep breathing in; (2) complete breathing out; (3) diaphragmatic aspiration (3 series of 8–12 repetitions). Breathing in was achieved by opening up the ribs, maximum breathing out by retracting the belly and completely releasing the air contained in the lungs, and diaphragmatic aspiration by acting as if one wanted to “suck one’s belly under the ribs” and maintain this position in apnea for 20 seconds. A standing, quadrupedic, and lying posture with active movements were selected, and the sequence of postures was left to the physiotherapist’s discretion. After diaphragmatic aspiration, a voluntary contraction of PFM was achieved and sustained throughout the active movements. A complete session lasted 30 *min*.

In ELEC group, a commercially available electrical stimulator (Myomed 632, Enraf Nonius, Delft, the Netherlands) was used to deliver transvaginal electrical stimulations (excitomotor, bidirectional, rectangular, symmetric current) via vaginal probes (Goode et al, 2003) and lasts 15 *min* (450 *s* on right PR and 450 *s* on left PR) and biofeedback training lasts 15 *min* (6 *s* muscle contraction and 12 *s* of rest). Frequency stimulation was set between 20 and 50 *Hz*. Biphasic pulses had duration of 1 *ms*. The current intensity was adjusted to the maximum level that could be tolerated comfortably, up to maximum 100 *mA*.

Written informed consent was obtained from all individual participants included in the study. No financial compensation was provided to participate to the study. Institutional review board approval is not required at our institution for MRI using standard pulse sequences. A Consolidated Standards of Reporting Trials (CONSORT) diagram showing enrollment, training allocation, and follow-up is presented in Fig. 1.

**Fig. 1.**
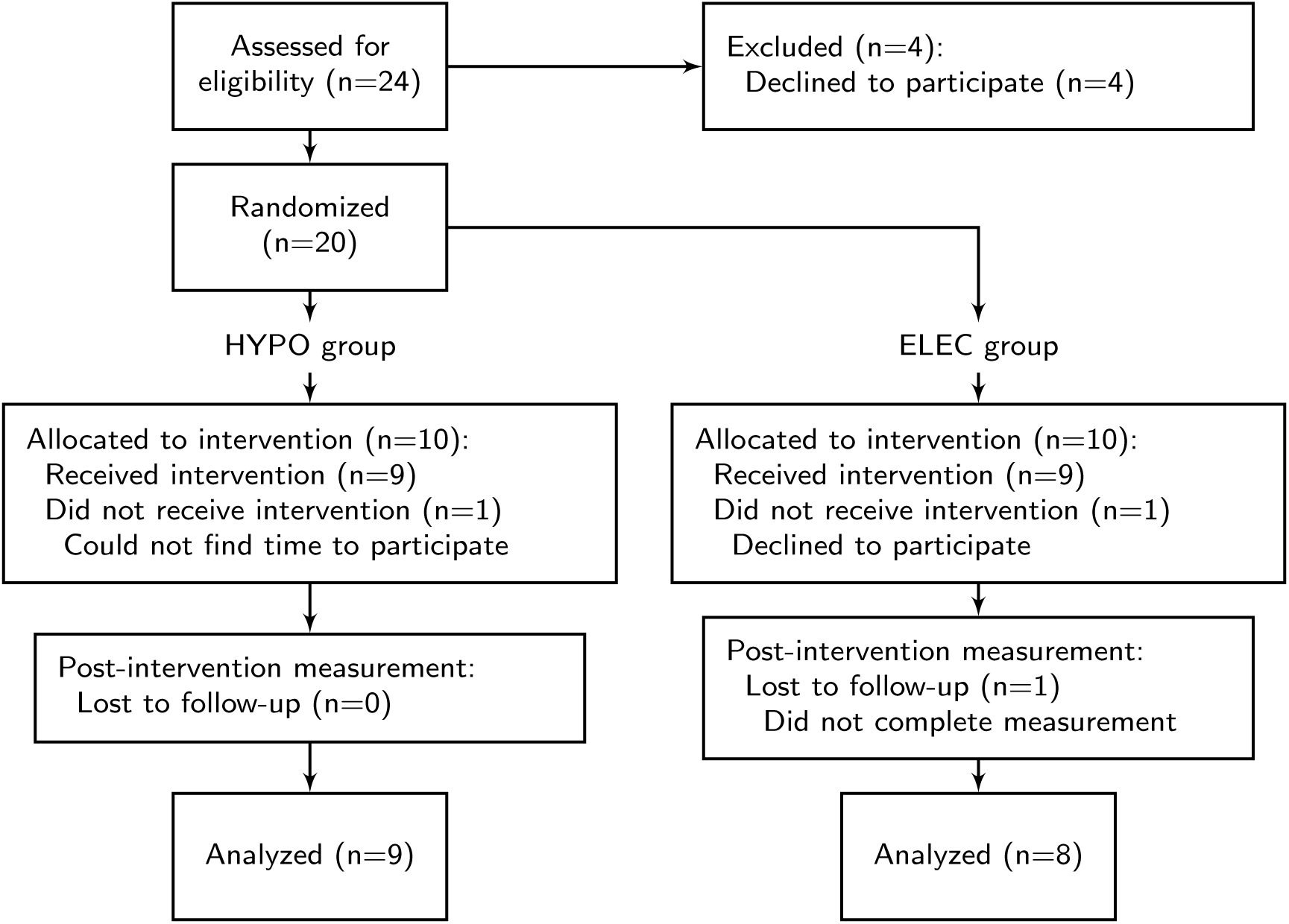
Flowchart of participants’ progress through the phases of the trial

### 2.2 Clinical and MRI assessments

Clinical assessment was realized in gynecological position (with the thighs flexed to 90 degrees) by a physiotherapist who has specialized in uro-gynecology; before and after PFMT. At the first assessment, all participants were instructed on how to correctly perform their PFM using vaginal palpation. The physiotherapist was blinded to group allocation.

PFM strength was measured through an arbitrary 0–12 points scale using a highly reliable (Isherwood and Rane, 2000) mechanical perineometer (PFX2, Cardio Design, Castle Hill, New South Wales, Australia) with a vaginal probe (26–28 *mm* diameter, active surface: 55 *mm* long, overall: 108 *mm* long). The cranial movement of the vaginal probe during measurement of PFM contraction was observed. Any contraction for which a retroversion of the hip or a Valsava maneuver was noticed were discounted. PFM strength was also assessed by bi-digital palpation (Bø and Sherburn, 2005) with the two distal phalanges inside the introitus vagina using the Modified Oxford Grading System (MOGS). It was individually assessed for right and left parts of PR and perineal body (PB), and consists of a 0–5 points scale: 0 = no contraction, 1 = flicker, 2 = weak, 3 = moderate, 4 = good (with lift) and 5 = strong. Three consecutive maximal contractions sustained for 5 seconds were recorded for perineometer and digital assessments, with a 30-second interval between efforts and the best of the 3 was registered.

PFM resting tone (RT) was quantified by digital palpation of left and right PR muscles (Dietz and Shek, 2008) on a 6-point scale with the following grades: 0 = muscle not palpable, 1 = muscle palpable but very flaccid, wide hiatus, minimal resistance to distension, 2 = hiatus wide but some resistance to distension, 3 = hiatus fairly narrow, fair resistance to palpation but easily distended, 4 = hiatus narrow, muscle can be distended but high resistance to distension, no pain, and 5 = hiatus very narrow, no distension possible, ‘woody’ feel, possibly with pain: ‘vaginismus’. PFM RT was computed as the sum of the grades of left and right parts of PR. Diaphragmatic aspiration (DA) (Caufriez, 1997) was rated on a scale of 0–5–10 (0 = no movement, 5 = mobilization of the perineum or viscera, 10 = mobilization of the perineum and viscera). The steps of DA are as follows: slow diaphragmatic inspiration, followed by total expiration and, after glottal closure, a gradual contraction of the abdominal wall muscles, with superior displacement of the diaphragm cupola (Caufriez, 1997; Resende et al, 2012; Santa Mina et al, 2015).

For MRI acquisition, each participant underwent an assessment performed in a supine position using a 1.5 Tesla MRI scanner (Ingenia 1.5T, Phillips, The Netherlands) and surface coils. Static T2-weighted turbo spin echo (TSE) techniques without fat saturation were used to image the sagittal, coronal, and axial planes during 20 minutes (spatial resolution 0.52×0.52 *mm*, repetition time (TR) 3745-4815 *ms*, echo time (TE) 90 *ms*, no gap, slices thickness 3 *mm* and water phase shift minimum). Seventy images were obtained in sagittal plane, fifty in coronal and thirty-five in the PR paraxial plane. All participants were instructed by a medical imaging technologist to relax their PFM while breathing in and out normally. MRI assessments were performed before and after PFMT. All images were obtained at our nuclear magnetic resonance department. Different measurements were performed on images and are detailed in Table 1, Fig. 2 and 3. Mid-pubic line (MPL) (Singh et al, 2001), pubococcygeal line (PCL) (Yang et al, 1991), and posterior levator plate (PLP) (Hsu et al, 2006) were determined (see Fig. 2 and Table 1).

**Fig. 2.**
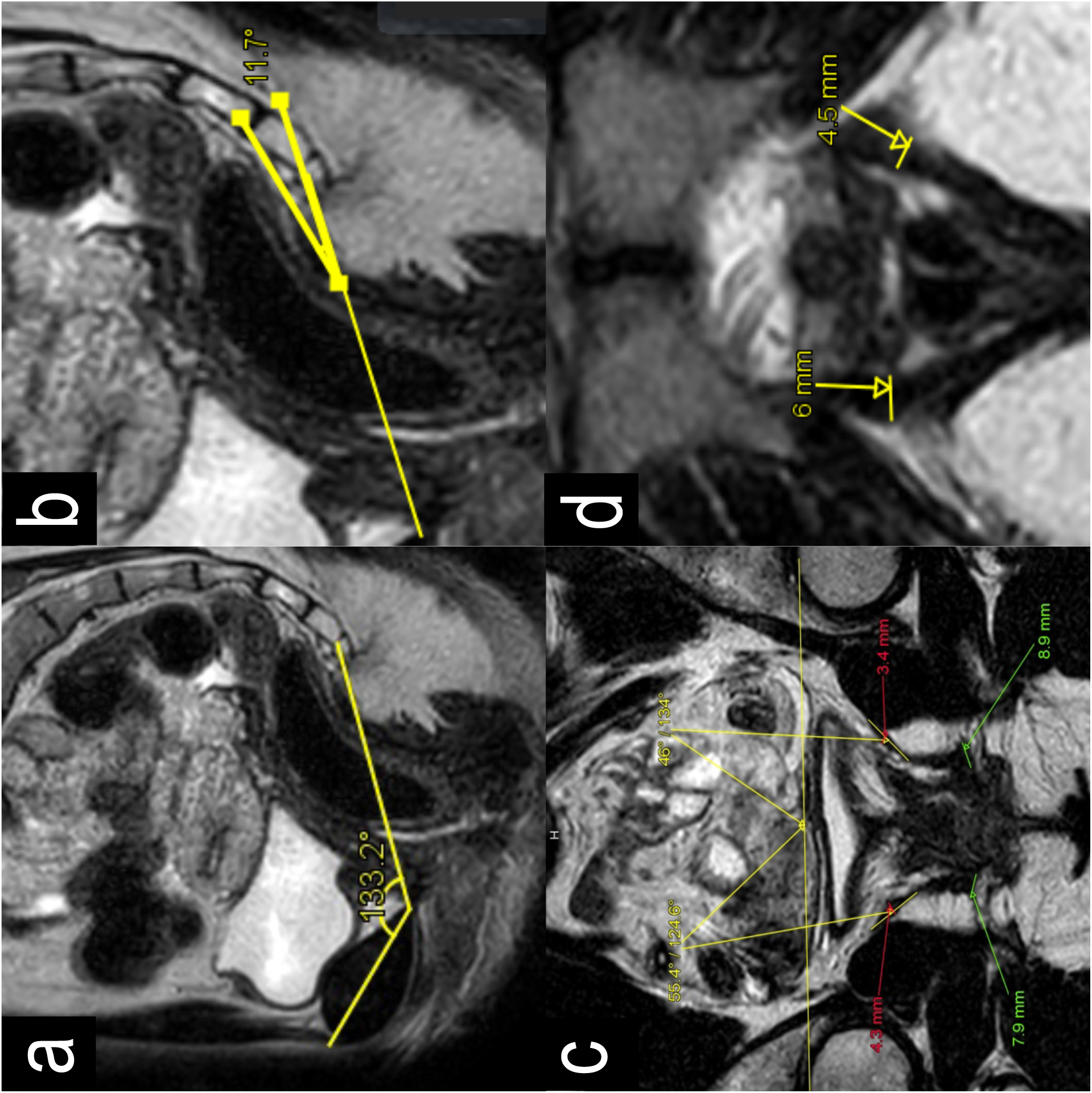
Different IRM views showing measurements. (a) midsagittal plane: MPL–PCL angle; (b) midsagittal plane: PLP–PCL angle; (c) coronal plane to the mid vagina: PR thickness (green), IL thickness(red) and angle (yellow); (d) para-axial view: PR thickness. Abbreviations: PR: *puborectalis*; IL: *iliococcygeus*; MPL: mid-pubic line; PCL: pubococcygeal line; PLP: posterior levator plate.

**Fig. 3.**
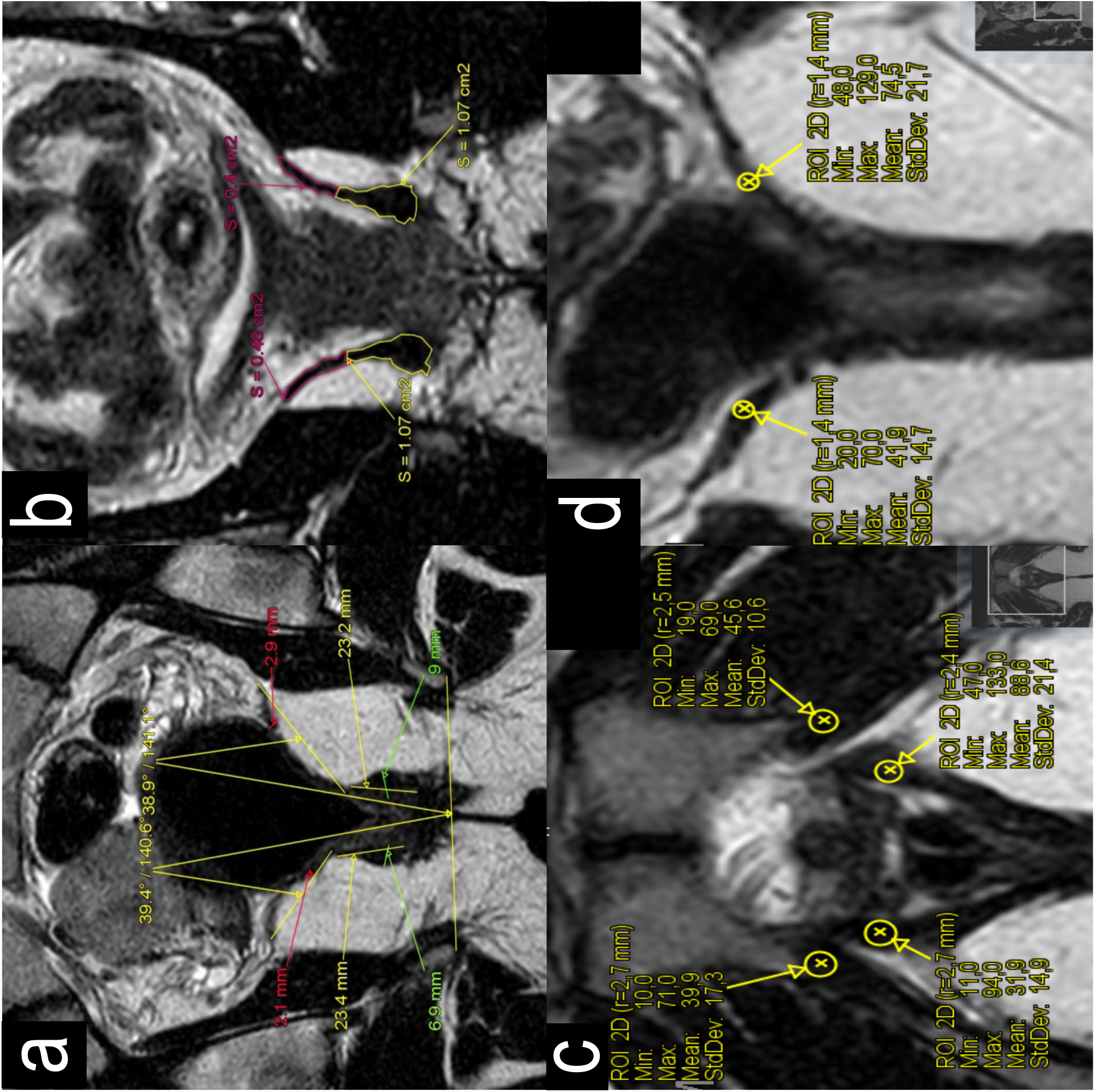
Different IRM views showing measurements. (a) coronal plane to the mid rectum: PR thickness (green) and height (yellow), IL thickness (red) and angle (yellow); (b) coronal plane: PR (yellow) and IL (red) contouring; (c) axial plane: PR signal intensity; (d) coronal plane: IL signals intensity. Abbreviations are the same as in Fig. 2.

**Table 1.** Measurements done on MRI. Illustrations are given in Fig. 2 and 3.

|  | Plane | Measurements<br>(units) | Measurement<br>sites | Sites<br>checked | Fig. # |
| --- | --- | --- | --- | --- | --- |
| **MPL–PCL**<br>**angle** | Midsagittal | Angle<br>(deg) | Between lower<br>edge of symphysis<br>pubis and last<br>sacrococcygeal disc<br>space | Through the<br>bladder, urethra,<br>vagina and rectum | 2a |
| **PLP–PCL**<br>**angle** | Midsagittal | Angle<br>(deg) | Between the point<br>of intersection<br>of levator plate-PCL and distal<br>insertion | Through the<br>bladder, urethra,<br>vagina and rectum | 2b |
| **PR**<br>**thickness** | Para-axial<br>(in PR axis,<br>at the level of<br>the inferior<br>border of the<br>pubic<br>symphysis) | Maximum<br>thickness<br>(mm) | Mid-rectum and<br>mid-vagina.<br>At 9 & 3 o'clock<br>the vagina | Sagittal | 2c<br>3a |
| **PR**<br>**height** | Coronal | Maximum<br>thickness<br>(mm) | Mid-rectum | Sagittal | 3a |
| **IL**<br>**thickness** | Coronal | Maximum<br>thickness<br>(mm) | Mid-rectum and<br>mid-vagina | Sagittal | 2c<br>3a |
| **IL**<br>**angle** | Coronal | Angle<br>(deg) | Mid-rectum and<br>mid-vagina.<br>Angle formed by IL<br>with horizontal<br>plane of pelvis | Sagittal | 2c<br>3a |
| **PR**<br>**volume** | Coronal | Contouring<br>(cm<sup>3</sup>) | All images were<br>the muscle appear | Sagittal | 3b |
| **IL**<br>**volume** | Coronal | Contouring<br>(cm<sup>3</sup>) | All images where<br>the muscle appear | Sagittal | 3b |
| **PR signal**<br>**intensity** | Axial | Relative signal<br>PR/OI×100<br>(%) | At 9 & 3 o'clock<br>the vagina and OI |  | 3c |
| **IL signal**<br>**intensity** | Coronal | Relative signal<br>IL/OI×100<br>(%) | Through back of<br>rectum |  | 3d |

PR: *puborectalis*; IL: *iliococcygeus*; OI: *obturator internus*; MPL: mid-pubic line; PCL: pubococcygeal line; PLP: posterior levator plate.

Measurements required about 6 hours for each participant and were performed by a radiologist who has specialized in pelvic floor MRI (L.M.) and two physiotherapists (E.G. and C.L.).

### 2.3 Statistical methods

All statistical procedures were performed with SigmaPlot software version 11.0 (Systat Software, San Jose, CA). The *p*-value was considerate significant when less than 0.05. Clinical results at baseline between HYPO and ELEC groups were compared using Mann-Whitney U tests and MRI results with *t*-tests. Wilcoxon signed-ranked tests were performed to examine the effects of training protocols on clinical results in HYPO and ELEC groups. A two-way RM ANOVA (before-after, training groups) with Holm-Sidak method for pairwise multiple comparisons was performed to examine the effect of PFMT and training protocols on MRI results. All data are presented as means and standard deviations (SD) and were checked for normality (Shapiro-Wilk) and equal variance (Brown-Forsythe) tests.

For PR and IL volumes, both interobserver and intraobserver reliability were assessed using intraclass correlation coefficient (ICC(2,k): 2-way random effects, absolute agreement, multiple raters (k=2): E.G. and L.M.; ICC(2,1): 2-way random effects, absolute agreement, single rater: E.G.) (Shrout and Fleiss, 1979). Each observer’s first measure recorded from each subject was used in the calculation of interobserver reliability. Intraobserver reliability was calculated using a second measurement recorded from each subject. ICCs were computed with R software (version 3.4.1) and irr package (version 0.84).

## 3 Results

### 3.1 Clinical results

None of the participants had PFD symptoms as determined by responses to questionnaires PFID-20 and PFQ-7. No significant differences between groups were observed at baseline for all clinical results. MOGS scores for left PR and PB significantly increased in the two groups but MOGS score for right PR significantly increased only in HYPO group (Table 2). PFM strength and resting tone, and diaphragmatic aspiration were not modified.

**Table 2.** Clinical results.

| *Training groups* | Before (B) |  | After (A) |  | B versus A |  |
| --- | --- | --- | --- | --- | --- | --- |
|  | HYPO | ELEC | HYPO | ELEC | W/*p* (HYPO) | W/*p* (ELEC) |
| **PFM strength** | 4.5[4-7.25] | 4[2.5-6] | 5[3-5.5] | 5.25[3.625-7.750] | 12/ 0.461 | 17/ 0.094 |
| **MOGS rPR** | 3[1.5-3.5] | 4[2-4] | 4[3.75-4.25] | 4[3.125-4.875] | 38/ **0.020** | 21/ 0.078 |
| **MOGS lPR** | 2.5[1.5-3.5] | 3[2-3.375] | 3.5[3-4.25] | 3.75[3-4.5] | 35/ **0.039** | 28/ **0.016** |
| **MOGS PB** | 3[1-4] | 4[1.25-4.375] | 4.5 [3.75-5] | 4[3.5-5] | 36/ **0.008** | 24/ **0.047** |
| **PFM RT** | 6[4.25-8] | 7.5[5.25-8.875] | 7[4.5-8] | 7[7-8] | 1/ 0.945 | 5/ 0.688 |
| **DA** | 10[5-10] | 7.5[5-10] | 10[5-10] | 7.5[5-10] | 0/ 1.000 | 0/ 1.000 |

PFM: pelvic floor muscles, MOGS: Modified Oxford Grading System, rPR: right *puborectalis*, lPR: left *puborectalis*, PB: perineal body, RT: resting tone, DA: diaphragmatic aspiration.

### 3.2 MRI results

No significant differences between groups were observed at baseline for all MRI results. The ICC values were 0.70 (inter-) and 0.95 (intra-) for PR volume and 0.80 and 0.83 for IL volume, showing moderate to very good reliability results. Muscle volumes of right and left IL significantly decreased after training and signal intensities of right and left PR significantly decreased after training (Table 3). Muscle thickness of right and left IL at mid-vagina location significantly decreased after training (Table 3). MPL–PCL and PLP–PCL angles, PR volume, IL signal intensities, and PR and IL measurements done at mid-vagina, mid-rectum, and at 9 & 3 o’clock the vagina were not modified.

**Table 3.**
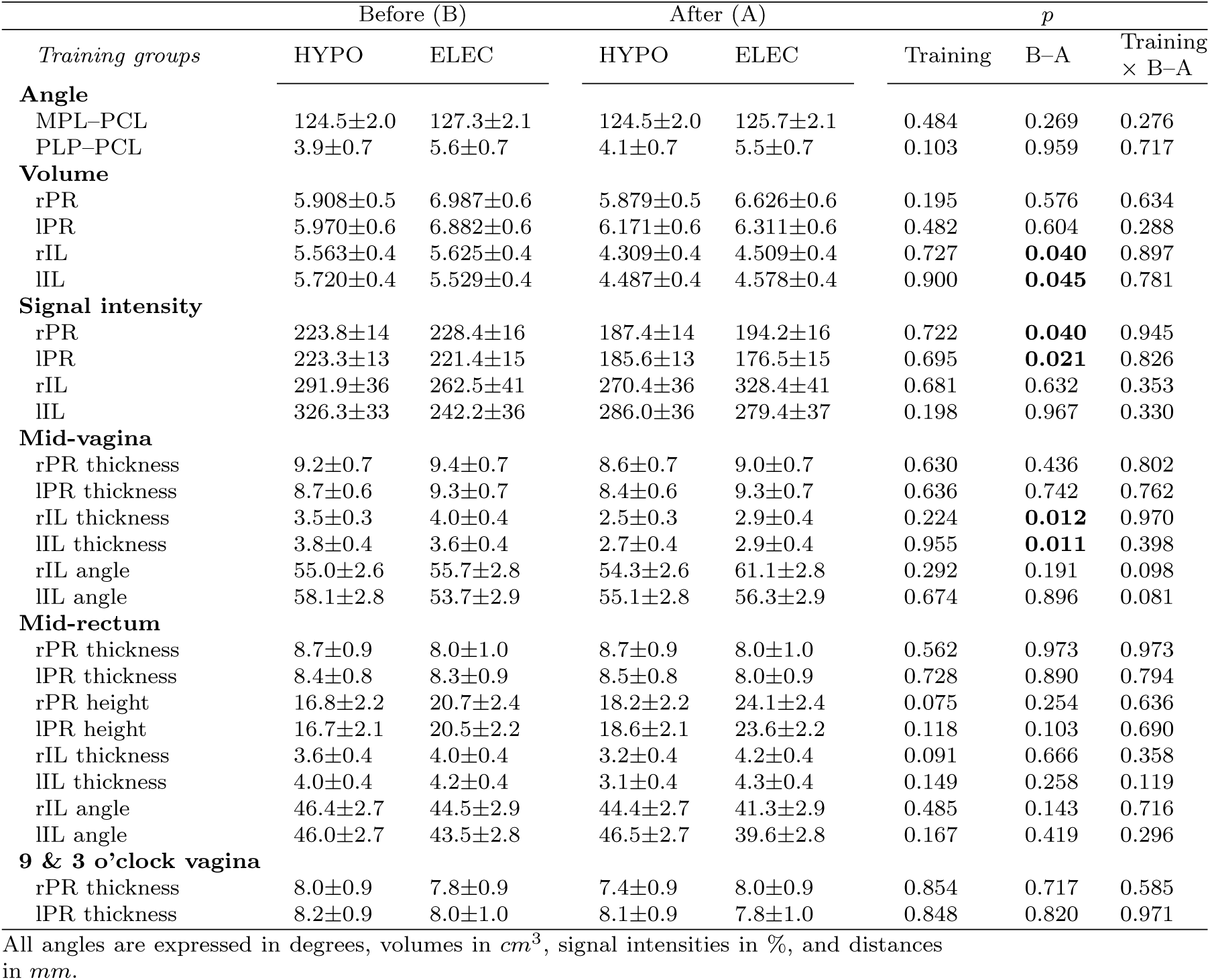
MRI results.

## 4 Discussion

LA muscle complex is one of the most important and complicated anatomical structure of the human body and as a result also one of the most poorly understood (Lammers et al, 2013). However, understanding the basic anatomy of the LA is essential when formulating a clinical opinion as injuries occur in 13–36% of women who have a vaginal delivery (Schwertner-Tiepelmann et al, 2012). Until now, no study explore the effects of a short period of intensive PFMT on the morphology of the pelvic muscles and related clinical scores in healthy nulliparous women, and the produced changes are uncertain. Our hypothesis was that the protocol based on biofeedback combined with vaginal stimulations would lead to less important morphological and clinical changes of PFM than voluntary contractions combined with hypopressive exercises. Results show the presence of clinical and morphological changes in different parts of LA muscle at rest after PFMT. However, changes were not significantly higher in the group that do not use instrumentation.

Historically, MRI studies have outlined the anatomy of PFM much more clearly than was possible with anatomical dissection studies (Raizada and Mittal, 2008), and different subdivisions of the LA were identified (Kearney et al, 2004; Margulies et al, 2006). *Terminologia Anatomica* (Federative Commitee on Anatomical Terminology, 1998), identifies three major subdivisions of LA: IL, PR, and *pubococcygeus* (PC). Over the years, these subdivisions have been given several names, making understanding even more difficult (Lammers et al, 2013). Among female, PC muscle, also called *pubovisceralis* (PV), is further divided into the *puboperinealis, pubovaginalis,* and *puboanalis* (Lawson, 1974; Kearney et al, 2004). Even if different parts of PV and enclosing urogenital hiatus, could be observed with standard MRI techniques, they can not be distinguished at their origin of the pubic bone (Zijta et al, 2013). Only MRI studies with diffusion tensor imaging (DTI) and fiber tractography allowed to provide a three-dimensional overall appearance of muscular fibers of PV (Zijta et al, 2011; Rousset et al, 2012), and it was therefore decided not to study PV.

From a morphological viewpoint, PR is thicker than IL and forms a band around the urethra, vagina and rectum (Singh et al, 2002). Our results show that PR thickness values before PFMT were greater at mid-vagina level compared to other levels, with mean values between 8.7±0.6 and 9.4±0.7 mm, and IL thickness values were greater at mid-rectum level, with mean values between 3.6±0.4 and 4.0±0.4 mm. Singh et al (2002) observed lower thickness values for PR (6.5±2.0) and IL (2.9±0.8), but their sample was composed of healthy nulliparous women aged between 23–42 *yrs.* Dumoulin et al (2007) showed in women with SUI that LA surface area at rest after PFMT was significantly smaller than before training; however, during a voluntary contraction, LA surface was significantly higher than before training. Even in healthy women, our results are in agreement with this study since we observed a decreased volume and thickness of IL muscle at mid-vagina level, both for PFMT with and without instrumentation. A dynamic MRI study is therefore needed to complete our static results. Furthermore, PR muscle volume and thickness were unchanged but signal intensity of PR compared with that of OI decreased, both for PFMT with and without instrumentation. The signal intensity of healthy muscles, *i.e*. muscles showing no muscle edema, fatty infiltration, or mass lesion, is well lower than that of fat for T2-weighted images (May et al, 2000). Hence it can be argued that the decreased signal intensity of PR indicates a decrease in fatty composition of this muscle, in favor of an increase of muscle fibers. Since SUI, apart of ligamentous and fascial lesions, is associated with increased signal intensity (Kirschner-Hermanns et al, 1993), this observation could be of major importance in pathological conditions. From a clinical viewpoint, MOGS scores significantly increased after PFMT, indicating a global increase of PFM strength. Since a 3-week period of intensive PFMT, based on 90 minutes/week, was sufficient to increase muscle strength in the 2 training groups, we believe that, in a preventive purpose, it is not necessary to use an expensive and more invasive method than combined voluntary and hypopressive exercises (Bernardes et al, 2012).

PFM strength changes observed after PFMT and assessed with perineometer were not significant. Perineometer values represent a global measurement of vaginal closure force that also includes the “noise” caused by the rise in intraabdominal pressure that often accompanies a LA contraction (Ashton-Miller et al, 2014). Even if we systematically checked the presence of a cranial movement of the vaginal probe during the measurement, we can exclude an increase in intra-abdominal pressure that could have masked the finding of a significant increase in the vaginal closure force after PFMT. The use of an instrumented speculum designed to minimize the effect of intra-abdominal pressure during measurement of PFM strength and developed by Ashton-Miller et al (2014), may be a solution to overcome this limitation. Another explanation of these results could be that the global maximal vaginal closure force was not modified but that some muscles have increased their strength to the detriment of others. Our hypothesis is based on the following observations after PFMT: (1) the decreasing volume and thickness of IL at the level of mid-vagina, without any change in signal intensity; and (2) the decrease of signal intensity in PR. Therefore, IL force may have been reduced while PR force increased. If this is the case, PFMT could be effective in modifying the resulting forces of the different parts of LA since the lines of action of IL (PV) and PR muscles are quite different: the first one lifts the pelvic floor while the second closes it (Betschart et al, 2014). The absence of a change in signal intensity of IL must be balanced according to the training groups. Surprisingly, signal intensity tends to increase in ELEC group while it decrease in HYPO. This finding also support the use of non-instrumented PFMT.

The present study has several limitations. The first limitation is the small number of the participants; therefore, our results may be not representative of the general population. To reduce bias about group homogeneity, several inclusion criteria were met. The first inclusion criterion was that all participants were young and nulliparous women. According to Slieker-ten Hove et al (2009), voluntary muscle contraction decrease with age, but there is no relation with parity. DeLancey et al (2003) found abnormalities in the LA muscle on MRI after vaginal delivery, that are not found in nulliparous. In women over age 60 years, the PFMs are significantly thinner both at rest and during contraction, compared to younger women (Bernstein, 1997). The second criterion was that participants were European women to avoid ethnic variability. Howard et al found a functional and morphological differences in the urethral sphincter and support system of nulliparous black and white women (Howard et al, 2000; Handa et al, 2008). The second limitation is that PFMs were only assessed with static MRI and a dynamic MRI study is needed to complete our results (Cai et al, 2013; Dumoulin et al, 2007). Others limitations relate to MRI acquisition. Each participant underwent MRI adapted at supine position with a mid-full bladder repletion. In the supine position, PFMs do not undergo organ weight and gravity (Fielding et al, 1998). A mid-full bladder repletion condition is difficult to obtain because of the ability of each bladder. During image analysis, we observed different degree of repletion of the bladder, which could influence our results. Finally, signal measurements were difficult to do coronally in IL (very thin) and in some PR and IL frequently fasciculated with fat striations.

## 5 Conclusions

This study suggests that 270 minutes of PFMT in healthy nulliparous women produces clinical and morphological changes in PFMs at rest. Muscle volume and thickness of IL at mid-vagina location decreased after training. PR volume was not modified but its signal intensity decreased, indicating a decrease in fatty composition of this muscle, in favor of an increase of muscle fibers. Additionally, MOGS scores increased after training, indicating that participants improved voluntary force contraction of their PFMs. Static MRI appears to be a relevant tool to highlight the impact of PFMT but is not sufficient to fully explain all clinical improvement and morphological changes. Further dynamic MRI studies and/or DTI-based tractography are needed to better understand the impact of training on the morphology of PFMs, and determine if these changes are sustainable and applicable to women with PFD.

## Acknowledgements

The authors would like to thank Caroline Lahaye and Jean-Louis Greffe for their assistance in the development of the protocol, and Jean-Claude Malherbe (Enraf-Nonius) for making the stimulation/biofeedback material available throughout the experiment. They also thank the financial support of Grand Hôpital de Charleroi (GHdC asbl), Philips SA, and the Haute Ecole Louvain en Hainaut for the MRI.

